# A high-quality genome assembly and annotation of the gray mangrove, *Avicennia marina*

**DOI:** 10.1101/2020.05.30.124800

**Authors:** Guillermo Friis, Joel Vizueta, David R. Nelson, Basel Khraiwesh, Enas Qudeimat, Kourosh Salehi-Ashtiani, Alejandra Ortega, Alyssa Marshell, Carlos M. Duarte, John A. Burt

## Abstract

The gray mangrove [*Avicennia marina* (Forsk.) Vierh.] is the most widely distributed mangrove species, ranging throughout the Indo-West Pacific. It presents remarkable levels of geographic variation both in phenotypic traits and habitat, often occupying extreme environments at the edges of its distribution. However, subspecific evolutionary relationships and adaptive mechanisms remain understudied, especially across populations of the West Indian Ocean. High-quality genomic resources accounting for such variability are also sparse. Here we report the first chromosome-level assembly of the genome of *A. marina*. We used a previously release draft assembly and proximity ligation libraries Chicago and Dovetail HiC for scaffolding, producing a 456,526,188 bp long genome. The largest 32 scaffolds (22.4 Mb to 10.5 Mb) accounted for 98 % of the genome assembly, with the remaining 2% distributed among much shorter 3,759 scaffolds (62.4 Kb to 1 Kb). We annotated 23,331 protein-coding genes using tissue-specific RNA-seq data, from which 13,312 were associated to GO terms. Genome assembly and annotated set of genes yield a 96.7% and 92.3% completeness score, respectively, when compared with the eudicots BUSCO dataset. Furthermore, an F_ST_ survey based on resequencing data successfully identified a set of candidate genes potentially involved in local adaptation, and revealed patterns of adaptive variability correlating with a temperature gradient in Arabian mangrove populations. Our *A. marina* genomic assembly provides a highly valuable resource for genome evolution analysis, as well as for identifying functional genes involved in adaptive processes and speciation.

## Introduction

Mangroves are a polyphyletic group of trees and shrubs that inhabit the intertidal zone of the tropic and sub-tropic coasts (Primavera et al. 2018; Hogarth 2015; Nagelkerken et al. 2008; Polidoro et al. 2010). Mangroves share several morphological and physiological adaptations to their harsh intertidal habitat, including traits for tolerance of high salinity, alternating dessication and submergence of soils across tidal cycles, and exposure of roots to hypoxic, sulfide-rich soils (Tomlinson 2016). Mangroves are of great ecological and economic importance, providing key functions such as high productivity, much of which is exported to surrounding ecosystems, and acting as breeding, nesting, nursery, foraging and shelter areas for a range of biota (Lee et al. 2014; Carugati et al. 2018; Primavera et al. 2018; Nagelkerken et al. 2008). Mangroves also serve as an important carbon sink, supporting climate change mitigation and adaptation potential (Duarte et al. 2013). The diversity of evolutionary origins and adaptive mechanisms found in mangroves provide compelling systems for studying patterns of trait evolution, lineage divergence and speciation (e.g., Zhou et al. 2007; Urashi et al. 2013; Xu et al. 2017b).

Of the approximately 70 mangrove species described, the gray mangrove *Avicennia marina* has the broadest latitudinal and longitudinal distribution (Tomlinson 2016; Spalding et al. 2010; Hogarth 2015). At least three sub-specific, partially allopatric taxa or ‘varieties’ have been described: *A. marina* var. *australasica*, restricted to southeastern Australia and New Zealand; *A. marina* var. *eucalyptifolia*, in northern regions of Australasia; and *A. marina* var. *marina*, that ranges from New Caledonia in the Pacific and across the entire Indian Ocean (Tomlinson 2016; Duke 1991; Li et al. 2016). The broad geographic distribution of *A. marina* is reflected in its presence across diverse environmental gradients (e.g. temperature, freshwater, sediment and nutrient supply, salinity, tidal range) and spatial settings (e.g. open coastlines, coastal lagoons, estuaries, deltas, coral fringes) (Duke 1990; Quisthoudt et al. 2012). Along with its widespread distribution, several aspects of *A. marina* biology make it a promising model organism among mangroves for the study of evolution based on genomic and molecular approaches. First, the broad environmental gradients present across the *A. marina* range are mirrored by remarkable geographic variation in phenotypic and life-history traits (Duke 1990; Tomlinson 2016), which makes them an suitable system for studying evolutionary processes related to dispersal, directional selection, and neutral evolution. Previous studies have reported the phylogenetic relationships for the sub-varieties of *A. marina* and congeneric species (Duke et al. 1998; Li et al. 2016; Nettel et al. 2008), yet the specific drivers of lineage diversification remain understudied. In particular, the extensive gray mangrove populations from the Eastern African and Arabian coasts have rarely been included in reported DNA sequence-based analyses, either for diversity screening or other purposes (e.g., Duke et al. 1998; Li et al. 2016). Second, *A. marina* can tolerate highly variable and extreme environmental conditions, and often occupies marginal, biologically limiting environments at the edges of its distribution (Morrisey et al. 2010). While several biological structures and mechanisms of the gray mangrove physiology have been described (Nguyen et al. 2015; Naidoo 2016), the genetic basis and pathways underlying such tolerance are still largely unknown (Mehta et al. 2005; Jithesh et al. 2006; Xu et al. 2017a). Understudied Arabian populations represent a particular gap due to their occurrence at the extremes of air and water temperature, salinity, and aridity (Ben-Hasan and Christensen 2019; Camp et al. 2018). Third, the *A. marina* genome is moderately small and structurally simple compared with other eukaryotes, and presents a limited amount of transposable elements (Lyu et al. 2018; Xu et al. 2017a; Das et al. 1995), which facilitates the identification of short polymorphisms and structural variants.

Previous studies on *A. marina* using high-throughput sequencing techniques and genomic approaches have been released. Two multi-species studies, including *A. marina*, explored patterns of convergent evolution at functional genes (Xu et al. 2017a) and transposable elements loads (Lyu et al. 2018), based on draft genomes that were recently made available online. Whole-genome assemblies of *A. marina* and several other mangrove taxa have recently been used for demographic inference (Guo et al. 2018) and convergent evolution analysis (He et al. 2020), but the underlying genomic data are not publicly accessible.

Here, we report a high-quality, chromosome-level assembly obtained using proximity ligation libraries as a resource for genome-based studies on *A. marina* and related mangrove species. An structural and functional annotation based on RNA-seq data from multiple tissues is also provided. In addition, we generated whole-genome shotgun data for a set of resequenced individuals from six different populations along the coasts of the Arabian Peninsula. We used these data to evaluate the performance of the assembly as a reference to study patterns of adaptive variability at the genomic level. Resequencing data is also used for organelle assembling.

## Materials and Methods

### Genome sequencing and assembly

A high-quality genome was produced combining the previously released draft assembly of *Avicennia marina* (Xu et al. 2017a; GenBank assembly accession: GCA_900003535.1, GCA_900174615.1; Lyu et al. 2018) and newly generated sequence data from proximity ligation libraries. Preparation of proximity ligation libraries Chicago and HiC as well as scaffolding with the software pipeline HiRise (Putnam et al. 2016) was conducted at Dovetail Genomics, LLC. The sequenced sample sequenced was leaf tissue obtained from an individual located at Ras Ghurab Island in the Arabian Gulf (Fig 1; Abu Dhabi, United Arab Emirates; 24.601°N, 4.566 °E). Briefly, for Chicago and the Dovetail HiC library preparation, chromatin was fixed with formaldehyde. Fixed chromatin was then digested with DpnII and free blunt ends were ligated. Crosslinks were reversed, and the DNA purified from protein, which was then sheared to ∼350 bp mean fragment size. Libraries were generated using NEBNext Ultra enzymes and Illumina-compatible adapters, and sequencing was carried out on an Illumina HiSeq X platform. Chicago and Dovetail HiC library reads were then used as input data for genome assembly for HiRise, a software pipeline designed specifically for using proximity ligation data to scaffold genome assemblies (Putnam et al. 2016). A previously reported draft genome of *Avicennia marina* (Lyu et al. 2018; Xu et al. 2017a; GenBank assembly accession: GCA_900003535.1) was used in the assembly pipeline, excluding scaffolds shorter than 1Kb since HiRise does not assemble them. Further details are provided in the Supplementary Information.

**Fig. 1.**
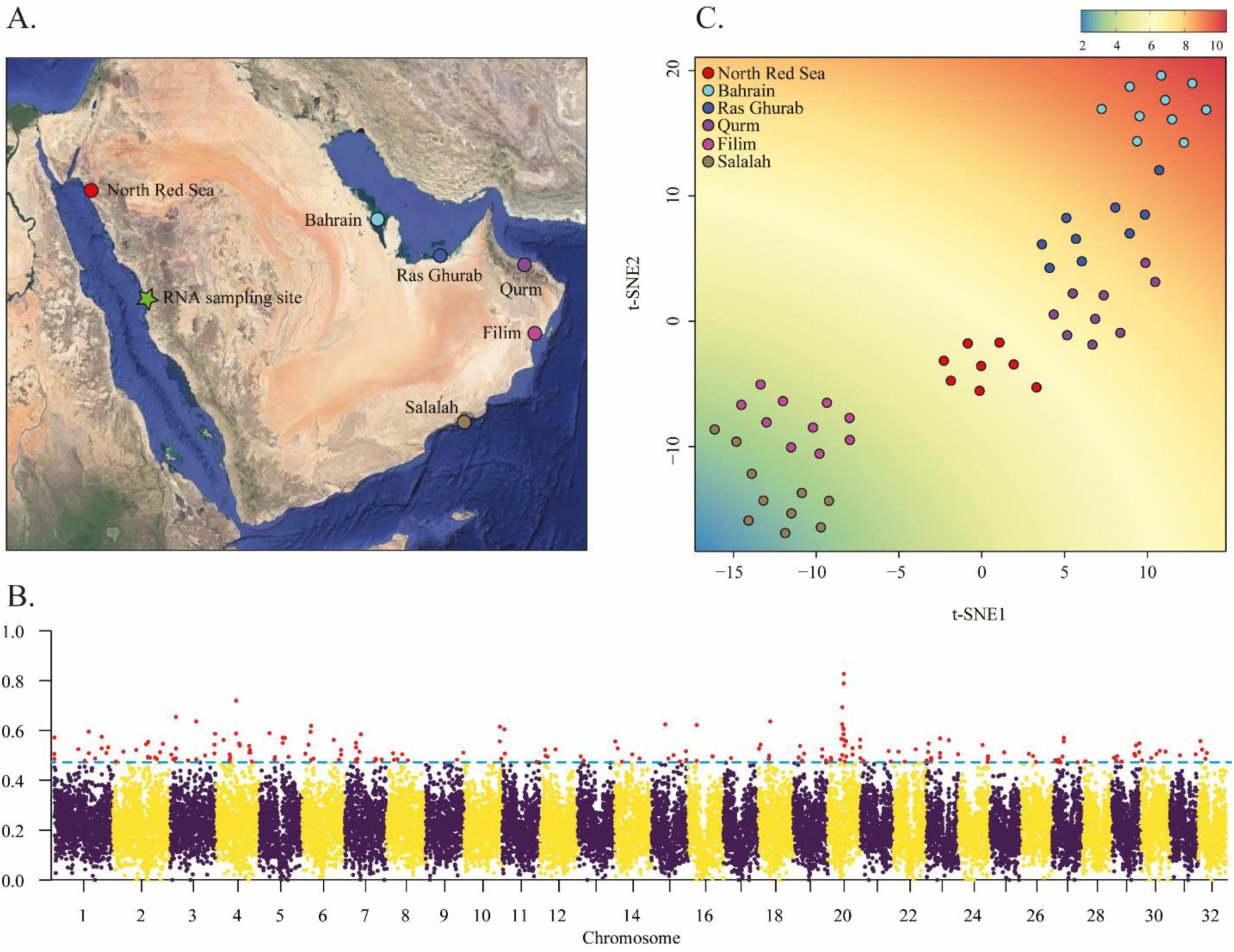
Geography and adaptive variability in Arabian gray mangroves. (A) Locations of the six stands sampled for whole genome resequencing (colored circles) and for RNA-seq (green star). (B) F_ST_ genome scan based on 22,181 windows of 20 Kb. Boxplot outliers (coefficient=1.5) are marked in red (C) t-SNE based on 200 SNP outliers linked to functional genes. The background shows the correlation between t-SNE1 and t-SNE2 with the annual temperature range registered in each one of the sampling locations. Temperature depicted in the legend is in °C.

The mitochondrial genome was assembled using NOVOplasty2.7.2 (Dierckxsens et al. 2017) and resequencing data based on Illumina paired-end 150 bp libraries from a conspecific individual (See below; Supplementary Information). The maturase (matR) mitochondrial gene available in NCBI (GenBank accession no. AY289666.1) was used for the input seed sequence. The assembly of the chloroplast yielded a high number of alternative, unsolved assemblies and is thereby no reported.

### Genome annotation

We performed the annotation of the *A. marina* genome using mRNA data from a set of tissues of conspecific individuals. Samples were collected on the coast of the Eastern Central Red Sea north of Jeddah in the Kingdom of Saudi Arabia (22.324 °N, 39.100 °E; Figure 1A). Total RNA was isolated from root, stem, leaf, flower, and seed using TRIzol reagent (Invitrogen, USA). RNA-seq libraries were prepared using TruSeq RNA sample prep kit (Illumina, Inc.), with inserts that range in size from approximately 100-400 bp. Library quality control and quantification were performed with a Bioanalyzer Chip DNA 1000 series II (Agilent), and sequenced in a HiSeq2000 platform (Illumina, Inc.). Messenger RNA reads were mapped with HISAT2 (Kim et al. 2015), and genome-referenced transcripts for each tissue were produced and merged with StringTie (Pertea et al. 2015). Prediction of coding regions was performed with TransDecoder (Haas and Papanicolaou 2015). The obtained gene annotation gff3 file was validated and used to generate the reported gene annotation statistics with GenomeTools (Gremme et al. 2013) and in-house Perl scripts. The Trinotate (Haas 2015) pipeline was then implemented to conduct a homology-based functional annotation by using Swiss-Prot (Bairoch and Apweiler 2000) and pfam (Bateman et al. 2002) databases, generating a final set of annotated functional genes. Further details on mRNA sequencing and annotation scripts are provided in the Supplementary Information.

Repetitive regions were first modelled *ab initio* using RepeatModeler v2.0.1 (Flynn et al. 2019) in all scaffolds longer than 100 Kbp with default options. The resulting repeat library was used to annotate and soft-mask repeats in the genome assembly with RepeatMasker 4.0.9 (Smit et al. 2015).

### Gene completeness assessment

We assessed gene completeness in the genome assembly, and gene annotation, using BUSCO (Benchmarking Universal Single-Copy Orthologs) v4.0.5 (--auto-lineage-euk option; Waterhouse et al. 2018). BUSCO evaluations were conducted using the 255 and 2,326 single-copy orthologous genes in Eukaryota_odb10 and Eudicots_odb10 datasets, respectively.

### *Adaptive variability analysis and functional assessment of* A. marina *genome*

To test the potential of the assembly and annotation reported here as a resource for genomic-based studies, we checked for regions of high divergence across the genome of *A. marina* using newly generated whole-genome data. We resequenced 60 individuals from six different populations from each of the major seas bordering Arabia (Figure 1A; Table S1, Supplementary Information), including populations in the Red Sea, the Arabian Gulf and the Arabian Sea/Sea of Oman. Arabia’s regional seas present extreme, but divergent, environmental conditions for mangrove growth. The northern Red Sea is characterized by cold winter temperatures and high salinity (Carvalho et al. 2019), while the southern Persian/Arabian Gulf is the world’s hottest sea each summer and is also hypersaline, with both areas considered arid to hyperarid with limited (<200 mm) rainfall (Vaughan et al. 2019). In contrast, the Arabian Sea and Sea of Oman have normal oceanic salinity, and summer temperatures that are buffered by cold-water upwelling as a result of the Indian Ocean monsoon, resulting in more benign environmental conditions (Claereboudt 2019). Using vcftools (Danecek et al. 2011), we conducted an F_ST_ survey across the six sampled populations based on 20 Kb sliding windows, and identifiied outlier loci associated with functional genes. We then used these loci to explore geographic patterns of adaptive variability by means of t-SNE analysis, testing for correlations between variability in sea surface temperature and t-SNE scores. Details on sequencing, variant calling, and analytical procedures are available in the Supplementary Information.

### Data availability

Genome assembly, genome annotation and related supporting resources for facilitating the use of these data have been deposited at DRYAD (doi:10.5061/dryad.3j9kd51f5). SNP matrix from resequenced individuals used in adaptive variation analyses is also provided in this dataset in vcf format. Raw resequencing data have been deposited in GenBank under the accessions SRA: SRP265707; BioProject: PRJNA629068.

The genome assembly has also been deposited at DDBJ/ENA/GenBank along with raw sequence data from Chicago and HiC libraries under the accession JABGBM000000000, SRA: SRP265707; BioProject: PRJNA629068.

Datasets relating to the RNA-seq analysis have been deposited in Mendeley (doi:10.17632/9tsp7fr28r). The RNA-seq reads have been deposited at Genbank under the Bioproject (SRA) accession: PRJNA591919; Biosample accession: SAMN13391520.

## Results and Discussion

### Sequencing and genome assembly

We sequenced and assembled a reference genome of a gray mangrove individual from the Arabian Gulf, an extreme environment at the northern limit of the species’ distribution. Chicago and Dovetail HiC libraries produced 235 million and 212 million 2×150 bp paired-end reads, respectively, and 134.1 Gb data overall. Genome scaffolding with HiRise yielded an assembly of 456.5 Mb for a final sequence coverage of 293X; an L50/N50 equal to 15 scaffolds/13.98 Mb and a relatively large number of ambiguous bases (i.e., N) inserted in the genome (10.6%; Table 1). The scaffold length distribution was heavily skewed towards extreme values (Table 1, Figure 2). The 32 longest scaffolds, ranging from 22.40 Mb to 10.58 Mb (median = 13.44 Mb) accounted for 98.03% of the whole assembled genome, congruent with a chromosome number 2N = 64 reported earlier (He et al. 2020). The remaining 1.97% of the genome was distributed among another 3,759 scaffolds ranging from 62.5 Kb to over 1 Kb (median = 1.8 Kb). The large number of small scaffolds may be due to the high fragmentation of the draft genome used in the assembly pipeline (Dovetails, personal communication). The integrity assessment of the *A. marina* genome retrieved a 98.8% and a 96.7% of the tested BUSCO groups for the eukaryote and the eudicots databases, respectively (Table 1). The remarkable discontinuity in length sizes, as well as the integrity and high quality of the scaffolding lends considerable support to the hypothesis of 32 chromosomes; further sequencing efforts involving long-read sequencing are warranted for confirmation. The mitochondrial genome assembly was 22,019 pb long with a 46.4% GC content.

**Table 1.** Summary statistics for the genome assembly and annotation of *A. marina*. CDS indicates protein-coding sequences.

| Genome assembly |  |
| --- | --- |
| Total length | 456,526,188 bp |
| Number of scaffolds | 3,791 |
| N50/L50 | 13,979,447 bp/15 scaffolds |
| N90/L90 | 11,144,373 bp/29 scaffolds |
| Chromosome scale | 10,583,658 bp/32 scaffolds |
| Longest scaffold | 22,400,447 bp |
| Missingness | 10.6% |
| GC content | 35.2 % |
| BUSCO eukaryota database | C:98.8% [S:81.2%, D:17.6%], F:0.8%, M:0.4%, N:255 |
| BUSCO eudicots database | C:96.7% [S:89.2%, D:7.5%], F:0.8%, M:2.5%, N:2,326 |
| Genome annotation |  |
| Number of genes | 23,331 |
| Number of annotated genes | 21,147 |
| Number of genes with GOs | 13,312 |
| Average gene length | 5,383.7 bp |
| Number of CDS | 59,445 |
| Average CDS length (bp) | 11,46.33 bp |
| Number of exons | 493,357 |
| Average exon length (bp) | 307.06 bp |
| Number of introns | 433,912 |
| Average intron length (bp) | 625.19 bp |
| BUSCO eukaryota database | C:97.2% [S:78.8%, D:18.4%], F:1.6%, M:1.2%, N:255 |
| BUSCO eudicots database | C:92.3% [S:85.2%, D:7.1%], F:1.6%, M:6.1%, N:2,326 |

BUSCO parameters are C: Complete BUSCO; S: Complete and single-copy BUSCOs; D: Complete and duplicated BUSCOs; F: Fragmented BUSCOs; M: Missing BUSCOs; and N: Total BUSCO groups searched. CDS indicates protein-coding sequences.

**Fig. 2.**
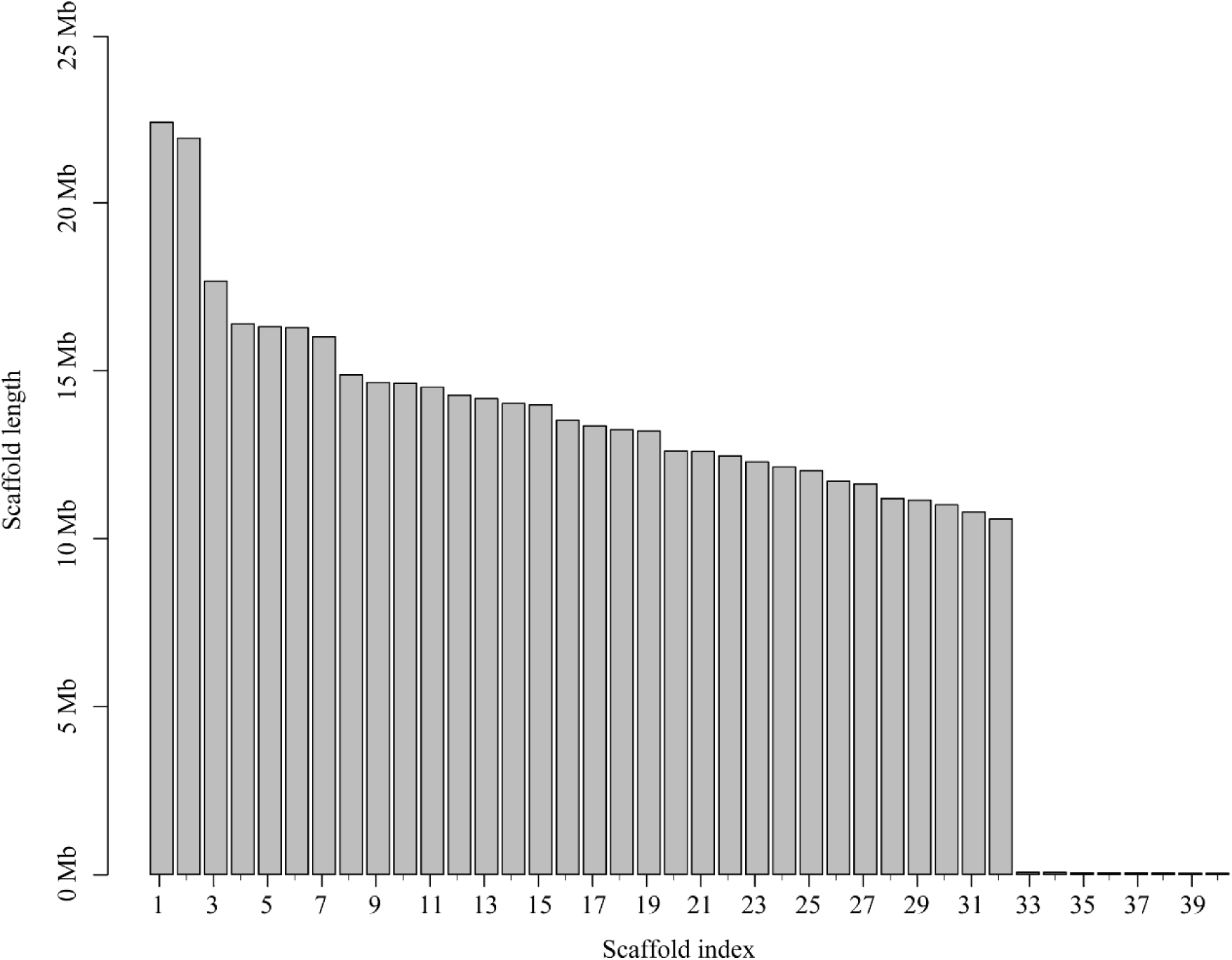
Length bar-plot of the longest 40 scaffolds arranged by decreasing size. The genome was sequenced using proximity ligation libraries Chicago and Dovetails HiC, and the assembly was carried out with HiRise pipeline.

### Genome annotation

We identified 23,331 protein-coding genes for which 21,147 orthologs from other species were identified, and 13,312 were associated to GO terms. The average gene length was 5.38 Kb, with a mean of 8.3 exons and 18.6 introns per gene. BUSCO integrity analysis reported a 97.2% of recovered BUSCOs for the eukaryota database, and a 92.3% in the case of the eudicots (Table 1). We also found that a total of 40.20% (188.5 Mb) of the *A. marina* assembly consisted of repetitive elements, a value moderately larger than the 30.4% previously reported for the species (Xu et al. 2017a). The greatest proportions corresponded to long terminal repeats and unclassified elements (20% and 16.7%, respectively; Table S2, Supplementary Information).

### *Adaptive variability analysis and functional assessment of* A. marina *genome*

We resequenced 60 individuals of *A. marina* from six different populations across the environmentally diverse coasts of the Arabian Peninsula (Figure 1A) with a coverage of 85X. After SNP calling and a strict filtering for quality and missing data, we obtained a dataset of 538,185 SNPs for 56 individuals. An F_ST_ scan based on sliding 20 Kb windows revealed a heterogeneous landscape of differentiation and detected a peak of high divergence at the Scaffold 20 (Figure 1B). A total of 200 highly divergent loci were identified, from which 43 overlapped with annotated genes associated to GO terms (Table S2, Figure S1). Several of these genes are involved in the development of shoots, leaves and flowers (BAM2 and BAM7; DeYoung et al. 2006; Reinhold et al. 2011), root and seeds (FEI1; Basu et al. 2016), as well as in protein storage (VTI12; Sanmartín et al. 2007). Importantly, we also found signals of differentiation in genes involved in plant sensitivity to salt and osmotic stress (WRKY40; Chen et al. 2010), drought and palatability to detritivorous crustaceans (LOX6; Grebner et al. 2013), supporting the role of abiotic and biotic factors in the differentiation of Arabian mangroves (Table S3, Supplementary Information). A t-SNE based 324 SNPs extracted from the functionally annotated, highly divergent loci showed clear clustering patterns among sampled populations (Figure 1C). Loading scores of retained t-SNE axes showed high correlation with the gradient of sea surface temperature (SST; p-values below 8.3×10^−14^ and 2.0×10^−16^ for t-SNE1 and t-SNE2, respectively), also congruent with a differentiation process driven by environmental factors. In light of these results, further research into gene evolution and biological pathways involved in local adaptation to the extreme environment of the Arabian mangroves is in process. These questions are, however, beyond the scope of this report and thus will be presented elsewhere.

In conclusion, we report the first chromosome-scale assembly for the *Avicennia marina* genome along with a comprehensive annotation based on tissue-specific RNA-seq data. The genome is highly contiguous and complete, and we demonstrated that it is a valuable resource for variant calling and the identification of functional, candidate genes underlying phenotypic and environmental divergence among mangrove taxa. Improved scaffolding also enables the identification of regions putatively under selection, including structural variants such as chromosome rearrangements or copy number variations, all relevant for investigating questions related to evolutionary biology and molecular ecology in this ecological and socioeconomically important species.

## Acknowledgements

The authors are grateful to NYUAD’s Core Bioinformatics for analysis and computational assistance. We thank Mark Priest, Dain McParland, Noura Al-Mansoori, Anique Ahmad, and Ada Kovaliukaite for their participation in the fieldwork. We gratefully acknowledge the Environment Agency Abu Dhabi (permit reference number 20181823a), and the Oman Ministry of Environment and Climate Affairs, Director General of Nature Conservation (permit number: 6210/10/75), for providing permits for research related to this manuscript. This work was supported by New York University Abu Dhabi Institute Grant (73 71210 CGSB9), NYUAD Faculty Research Funds (AD060) and King Abdullah University of Science and Technology baseline funding to CMD. We dedicate this manuscript to the memory of Basel Khraiwesh, who had produced the RNA-seq data used for annotation in this study.

## References

Bairoch, A., and R. Apweiler, 2000 The SWISS-PROT protein sequence database and its supplement TrEMBL in 2000. Nucleic Acids Research 28 (1):45–48.

Basu, D., L. Tian, T. Debrosse, E. Poirier, K. Emch et al., 2016 Glycosylation of a fasciclin-like arabinogalactan-protein (SOS5) mediates root growth and seed mucilage adherence via a cell wall receptor-like kinase (FEI1/FEI2) pathway in Arabidopsis. PLOS One 11 (1).

Bateman, A., E. Birney, L. Cerruti, R. Durbin, L. Etwiller et al., 2002 The Pfam protein families database. Nucleic Acids Research 30 (1):276–280.

Ben-Hasan, A., and V. Christensen, 2019 Vulnerability of the marine ecosystem to climate change impacts in the Arabian Gulf—an urgent need for more research. Global ecology and conservation 17:e00556.

Camp, E.F., V. Schoepf, P.J. Mumby, L.A. Hardtke, R. Rodolfo-Metalpa et al., 2018 The future of coral reefs subject to rapid climate change: lessons from natural extreme environments. Frontiers in Marine Science 5:4.

Carugati, L., B. Gatto, E. Rastelli, M.L. Martire, C. Coral et al., 2018 Impact of mangrove forests degradation on biodiversity and ecosystem functioning. Scientific Reports 8 (1):13298.

Carvalho, S., B. Kürten, G. Krokos, I. Hoteit, and J. Ellis, 2019 The Red Sea, pp. 49–74 in World Seas: An Environmental Evaluation, edited by C. Sheppard. Elsevier.

Chen, H., Z. Lai, J. Shi, Y. Xiao, Z. Chen et al., 2010 Roles of Arabidopsis WRKY18, WRKY40 and WRKY60 transcription factors in plant responses to abscisic acid and abiotic stress. BMC plant biology 10 (1):281.

Claereboudt, M.R., 2019 Oman, pp. 25–48 in World Seas: An Environmental Evaluation, edited by C. Sheppard. Elsevier.

Danecek, P., A. Auton, G. Abecasis, C.A. Albers, E. Banks et al., 2011 The variant call format and VCFtools. Bioinformatics 27 (15):2156–2158.

Das, A.B., U.C. Basak, and P. Das, 1995 Variation in nuclear DNA content and karyotype analysis in three species of Avicennia, a tree mangrove of coastal Orissa. Cytobios 84:93–102.

DeYoung, B.J., K.L. Bickle, K.J. Schrage, P. Muskett, K. Patel et al., 2006 The CLAVATA1-related BAM1, BAM2 and BAM3 receptor kinase-like proteins are required for meristem function in Arabidopsis. The Plant Journal 45 (1):1–16.

Dierckxsens, N., P. Mardulyn, and G. Smits, 2017 NOVOPlasty: de novo assembly of organelle genomes from whole genome data. Nucleic Acids Research 45 (4):e18–e18.

Duarte, C.M., I.J. Losada, I.E. Hendriks, I. Mazarrasa, and N. Marbà, 2013 The role of coastal plant communities for climate change mitigation and adaptation. Nature Climate Change 3 (11):961–968.

Duke, N., 1990 Phenological trends with latitude in the mangrove tree Avicennia marina. The Journal of Ecology:113–133.

Duke, N., 1991 A systematic revision of the mangrove genus Avicennia (Avicenniaceae) in Australasia. Australian Systematic Botany 4 (2):299–324.

Duke, N.C., J.A. Benzie, J.A. Goodall, and E.R. Ballment, 1998 Genetic structure and evolution of species in the mangrove genus Avicennia (Avicenniaceae) in the Indo-West Pacific. Evolution 52 (6):1612–1626.

Flynn, J.M., R. Hubley, C. Goubert, J. Rosen, A.G. Clark et al., 2019 RepeatModeler2: automated genomic discovery of transposable element families. bioRxiv:856591.

Grebner, W., N.E. Stingl, A. Oenel, M.J. Mueller, and S. Berger, 2013 Lipoxygenase6-dependent oxylipin synthesis in roots is required for abiotic and biotic stress resistance of Arabidopsis. Plant Physiology 161 (4):2159–2170.

Gremme, G., S. Steinbiss, and S. Kurtz, 2013 GenomeTools: a comprehensive software library for efficient processing of structured genome annotations. IEEE/ACM Transactions on Computational Biology and Bioinformatics 10 (3):645–656.

Guo, Z., X. Li, Z. He, Y. Yang, W. Wang et al., 2018 Extremely low genetic diversity across mangrove taxa reflects past sea level changes and hints at poor future responses. Global Change Biology 24 (4):1741–1748.

Haas, B., 2015 Trinotate: Transcriptome functional annotation and analysis.

Haas, B., and A. Papanicolaou, 2015 TransDecoder (find coding regions within transcripts). Github, nd https://github.com/TransDecoder/TransDecoder (accessed May 17, 2018).

He, Z., S. Xu, Z. Zhang, W. Guo, H. Lyu et al., 2020 Convergent adaptation of the genomes of woody plants at the land-sea interface. National Science Review.

Hogarth, P.J., 2015 The biology of mangroves and seagrasses: Oxford University Press.

Jithesh, M., S. Prashanth, K. Sivaprakash, and A. Parida, 2006 Monitoring expression profiles of antioxidant genes to salinity, iron, oxidative, light and hyperosmotic stresses in the highly salt tolerant grey mangrove, Avicennia marina (Forsk.) Vierh. by mRNA analysis. Plant cell reports 25 (8):865–876.

Kim, D., B. Langmead, and S. Salzberg, 2015 hisat2. Nature methods.

Lee, S.Y., J.H. Primavera, F. Dahdouh-Guebas, K. McKee, J.O. Bosire et al., 2014 Ecological role and services of tropical mangrove ecosystems: a reassessment. Global Ecology and Biogeography 23 (7):726–743.

Li, X., N.C. Duke, Y. Yang, L. Huang, Y. Zhu et al., 2016 Re-evaluation of phylogenetic relationships among species of the mangrove genus Avicennia from Indo-West Pacific based on multilocus analyses. PLOS One 11 (10):e0164453.

Lyu, H., Z. He, C.I. Wu, and S. Shi, 2018 Convergent adaptive evolution in marginal environments: unloading transposable elements as a common strategy among mangrove genomes. New phytologist 217 (1):428–438.

Mehta, P.A., K. Sivaprakash, M. Parani, G. Venkataraman, and A.K. Parida, 2005 Generation and analysis of expressed sequence tags from the salt-tolerant mangrove species Avicennia marina (Forsk) Vierh. Theoretical and Applied Genetics 110 (3):416–424.

Morrisey, D.J., A. Swales, S. Dittmann, M.A. Morrison, C.E. Lovelock et al., 2010 The ecology and management of temperate mangroves, pp. 43–160. CRC Press, Boca Raton(USA).

Nagelkerken, I., S. Blaber, S. Bouillon, P. Green, M. Haywood et al., 2008 The habitat function of mangroves for terrestrial and marine fauna: a review. Aquatic Botany 89 (2):155–185.

Naidoo, G., 2016 The mangroves of South Africa: An ecophysiological review. South African Journal of Botany 107:101–113.

Nettel, A., R.S. Dodd, Z. Afzal-Rafii, and C. Tovilla-Hernández, 2008 Genetic diversity enhanced by ancient introgression and secondary contact in East Pacific black mangroves. Molecular Ecology 17 (11):2680–2690.

Nguyen, H.T., D.E. Stanton, N. Schmitz, G.D. Farquhar, and M.C. Ball, 2015 Growth responses of the mangrove Avicennia marina to salinity: development and function of shoot hydraulic systems require saline conditions. Annals of Botany 115 (3):397–407.

Pertea, M., G.M. Pertea, C.M. Antonescu, T.-C. Chang, J.T. Mendell et al., 2015 StringTie enables improved reconstruction of a transcriptome from RNA-seq reads. Nature Biotechnology 33 (3):290.

Polidoro, B.A., K.E. Carpenter, L. Collins, N.C. Duke, A.M. Ellison et al., 2010 The loss of species: mangrove extinction risk and geographic areas of global concern. PLOS One 5 (4):e10095.

Primavera, J.H., D.A. Friess, H. Van Lavieren, and S.Y. Lee, 2018 The Mangrove Ecosystem, pp. 18–51 in World Seas: An Environmental Evaluation: Volume II: The Indian Ocean to the Pacific, edited by C. Sheppard. Academic Press.

Putnam, N.H., B.L. O’Connell, J.C. Stites, B.J. Rice, M. Blanchette et al., 2016 Chromosome-scale shotgun assembly using an in vitro method for long-range linkage. Genome Research 26 (3):342–350.

Quisthoudt, K., N. Schmitz, C.F. Randin, F. Dahdouh-Guebas, E.M. Robert et al., 2012 Temperature variation among mangrove latitudinal range limits worldwide. Trees 26 (6):1919–1931.

Reinhold, H., S. Soyk, K. Šimková, C. Hostettler, J. Marafino et al., 2011 β-Amylase–Like proteins function as transcription factors in Arabidopsis, controlling shoot growth and development. The Plant Cell 23 (4):1391–1403.

Sanmartín, M., A. Ordóñez, E.J. Sohn, S. Robert, J.J. Sánchez-Serrano et al., 2007 Divergent functions of VTI12 and VTI11 in trafficking to storage and lytic vacuoles in Arabidopsis. Proceedings of the National Academy of Sciences 104 (9):3645–3650.

Smit, A., R. Hubley, and P. Green, 2015 RepeatMasker Open-4.0. 2013–2015.

Spalding, M., M. Kainuma, and L. Collins, 2010 World atlas of mangroves. London, UK: Earthscan.

Tomlinson, P.B., 2016 The botany of mangroves: Cambridge University Press.

Urashi, C., K.M. Teshima, S. Minobe, O. Koizumi, and N. Inomata, 2013 Inferences of evolutionary history of a widely distributed mangrove species, Bruguiera gymnorrhiza, in the Indo-West Pacific region. Ecology and Evolution 3 (7):2251–2261.

Vaughan, G.O., N. Al-Mansoori, and J.A. Burt, 2019 The Arabian Gulf, pp. 1–23 in World Seas: An Environmental Evaluation, edited by C. Sheppard. Elsevier.

Waterhouse, R.M., M. Seppey, F.A. Simão, M. Manni, P. Ioannidis et al., 2018 BUSCO applications from quality assessments to gene prediction and phylogenomics. Molecular Biology and Evolution 35 (3):543–548.

Xu, S., Z. He, Z. Guo, Z. Zhang, G.J. Wyckoff et al., 2017a Genome-wide convergence during evolution of mangroves from woody plants. Molecular Biology and Evolution 34 (4):1008–1015.

Xu, S., Z. He, Z. Zhang, Z. Guo, W. Guo et al., 2017b The origin, diversification and adaptation of a major mangrove clade (Rhizophoreae) revealed by whole-genome sequencing. National Science Review 4 (5):721–734.

Zhou, R.C., K. Zeng, W. Wu, X.S. Chen, Z.H. Yang et al., 2007 Population genetics of speciation in nonmodel organisms: I. Ancestral polymorphism in mangroves. Molecular Biology and Evolution 24 (12):2746–2754.

